# Evolutionary Adaptation of Prephenate Dehydrogenases: A regulatory ACT domain acquisition in ecological niche specialization

**DOI:** 10.1101/2025.10.30.685691

**Authors:** A. Gritsunov, I.G. Shabalin, J. Hou, W. Minor, D. Christendat

## Abstract

Bacteria prephenate dehydrogenase (PDH) participates in the metabolic pathway for tyrosine biosynthesis. PDHs within the *Bacilaceae* phylum contain an ACT domain which enables them to be allosterically regulated by tyrosine. The mechanism via which the ACT domain introduces allostery onto PDH enzymes remains elusive. Furthermore, the evolutionary and biological advantages of ACT domain mediated regulation of metabolic pathways are highly debated. Building on our previous study, in which we solved the crystal structure of a *Bacillus antraces* ACT-containing PDH and proposed a model for its allosteric regulation by tyrosine, we now present further structural, and functional analyses in support of this model. In this study, we generated truncated PDH protein constructs lacking the ACT domain, determined their crystal structure and evaluated the role of tyrosine in modulating their enzymatic activity. We determined that the truncated PDH remains catalytically active, however, it is no longer allosterically regulated by tyrosine. Comparative structural analysis between the truncated PDH and PDHs naturally lacking the ACT domain that are known to be competitively inhibited by tyrosine revealed only minor changes in a loop region in the prephenate binding site. Attempts to introduce amino acids identified from the competitively inhibited PDH into the truncated construct did not restore tyrosine sensitivity, even at high concentration. This indicates that additional main chain amino acids away from the substrate binding site also contribute competitive inhibition by tyrosine. Analysis of a highly represented phylogenetic tree revealed that ACT containing PDHs are predominantly distributed amongst *Firmicute* and *Actinomycetota*. Representative organisms from both groups colonize nutrient limited and extreme environments. This distribution suggests that acquisition of the ACT domain may confer an evolutionary advantage by enabling organisms to efficiently partition chorismate, the end product of the shikimate pathway, for the biosynthesis of tyrosine and other essential aromatic compounds.

## Introduction

Tyrosine biosynthesis begins with the Claisen rearrangement of chorismate to prephenate, followed by either oxidative decarboxylation to hydroxyphenyl pyruvate or conversion to arogenate by an aminotransferase (Hagino and Nakayama 1974, Xia and Jensen 1990, Wiest and Houk 1994, Bonner, Jensen et al. 2004, Bonvin, Aponte et al. 2006, Sun, Singh et al. 2006). The enzymatic conversion of prephenate to hydroxyphenylpyruvate or arogenate represents the committed step in tyrosine biosynthesis and is therefore considered the primary regulatory point in the pathway. The enzymes belonging to the TyrA family catalyze these transformations of prephenate. They are classified based on their substrate preferences; prephenate dehydrogenase (PDH) utilizes prephenate, arogenate dehydrogenase utilizes arogenate, and cyclohexadienyl dehydrogenase can utilize either prephenate or arogenate as their substrates. All PDHs have a conserved catalytic core domain and a subgroup within this family also contain an ACT domain (Song, Bonner et al. 2005). In PDHs lacking the ACT domain, structural and biochemical studies have shown that the core catalytic domain is sufficient for both the enzymatic activity and regulatory properties. In this subgroup of ACT-lacking PDHs, tyrosine competes with respect to prephenate and inhibits PDH enzymatic activity (Sun, Shahinas et al. 2009).

In contrast, ACT containing prephenate dehydrogenases are regulated through an allosteric mechanism in which the ACT domain serves as the site for tyrosine binding and regulation (Shabalin, Gritsunov et al. 2020). The ACT domain is predominantly found in low-GC gram-positive bacteria (Song, Bonner et al. 2005), particularly in PDH-ACT protein that exhibit specificity for the NAD^+^ cofactor. Recent structural and biochemical data have provided insights into the allosteric mode of inhibition of ACT containing prephenate dehydrogenases (Shabalin, Gritsunov et al. 2020). The crystal structure of *B. antraces* ACT-containing PDH revealed dimerization of the core catalytic domain similar to non-ACT containing PDHs (Sun, Shahinas et al. 2009). The structure of the active site indicates that catalytic and cofactor binding residues are conserved between with members of PDH family. From a structural perspective, the symmetry within dimers is disrupted by the presence of the dimerized ACT domains, which sit atop of the catalytic cores. Within the crystal structure, the ACT domain is shifted towards one side of the dimer. This disruption in symmetry resulted in one ACT monomer becoming constricted and the other slightly extended in the dimer. The flexible loop connecting the ACT domain and dimerization domain protrudes into the active site. This structural effect is expected to interfere with ligand binding which serves a primary mechanistic role in ACT-PDH allostery. Enzyme kinetic further support this structural model in which increasing concentration of tyrosine directly reduce the maximal velocity of the catalyzed reaction of ACT-PDH as expected for classical allosteric inhibition.

The crystal structure of BaPDH-ACT provides a solid framework for investigating the regulatory mechanism of tyrosine allosteric inhibition. It enables us to investigate of the role of key regulatory structural elements of the protein. However, the absence of a tyrosine-free-structure limits greater understanding of the conformational changes associated with allosteric inhibition. To address this BaPDH was co-crystallized with tyrosine to elucidate its role in the allosteric inhibition of the protein. The structure of the tyrosine-BaPDH complex revealed that tyrosine binds to the ACT domain which we speculated induced distinct conformational change s to block the active site of the protein. A proposed mechanism for allosteric inhibition of BaPDH by tyrosine has already been described (Shabalin, Gritsunov et al. 2020). This study adds critical experimental mechanistic information to this mode of allosteric regulation. To do this we we prepared protein expression constructs lacking the ACT domain and in-parallel design site directed mutants to investigate the molecular mechanism involving tyrosine binding. By solving the crystal structure of the ACT truncated construct we provided mechanistic insight on tyrosine induced conformational changes that results in allosteric inhibition of BaPDH activity. In the absence of the ACT domain, symmetrical positioning of each monomer in the BaPDH dimer in complexed with NAD^+^. Similar placement of amino acid residues in the cofactor binding site and active site was observed as in the ACT containing version of the protein. The positioning of the Lasso loops in the truncated construct above the cofactor site were found to be in the open conformation, thus allowing unrestricted access of NAD^+^ and prephenate to the active site of the protein. This is consistent with kinetic analysis, demonstrating that the activity of the truncated construct does not undergo allosteric inhibition with tyrosine.

## Methods

### Mutagenesis and truncation of BaPDH construct

The Bacillus anthracis PDH construct from pMCSG7 vector (Shabalin, Gritsunov et al. 2020), BaPDH, containing an N-terminal 6 × His affinity tag with a TEV protease recognition cleavage site was used as a template for generating site-directed mutants and the ACT domain deletion construct. Site-directed mutation was prepared using the quick change mutagenesis approach described elsewhere (Liu and Naismith 2008). The BaPDH ACT deletion construct (306-378)Δ was prepared by sub cloning the respective DNA region into pET-28mod plasmid (Singh and Christendat 2006). All mutants and constructs were verified by Sanger DNA sequencing.

### Expression and purification of BaPDH mutants

Protein expression and purification were carried out according to the established protocol for BaPDH purification (Shabalin, Gritsunov et al. 2020). Briefly, plasmids for the respective BaPDH variants were transformed into E. coli strain BL21-CodonPlus® competent cells and selected with either ampicillin or kanamycin antibiotics on the LB-Agar plate. Freshly transformed *E. coli* cells were inoculated into LB (Luria Broth) media with either 50 µg/ml kanamycin or 100 µg/ml ampicillin. Expression of the protein was induced by the addition of IPTG. To 0.4mM Cell lysis was conducted with sonication and French press at 1000psi. Lysates were spun down at 20,000xg for 30minutes; proteins were purified from the supernatant with nickel nitrilotriacetic acid affinity chromatography (NiNTA). The 6xHIS tag was cleaved from the purified protein with TEV protease incubation overnight at 4 °C and cleavage was monitored on a 12% SDS-acrylamide gel. The protein was then separated from the cleaved 6xHIS tag by passage through a nickel nitrilotriacetic acid column and dialyzed at 4°C in a buffer containing 50 mM Tris pH 7.5, 500 mM NaCl, and 5% glycerol. Dialyzed protein was concentrated to ∼10mg/ml, flash frozen with liquid nitrogen and stored at -80 C°. Protein purity was assessed by running 14% SDS PAGE.

### Enzyme kinetics

The enzymatic activity of BaPDH was assayed spectrophotometrically by monitoring the production of NADH at 340 nm (extinction coefficient 6,220 M^−1^cm^−1^) as previously described (Shabalin, Gritsunov et al. 2020). The reaction buffer consisted of 50 mM Tris pH 7.5, 150 mM NaCl, 2.5 mM MgCl2, 2 mM DTT. Appropriate concentrations of NAD^+^ and prephenate were added to initiate the reaction. Enzyme kinetics reactions were conducted using a fixed saturating concentration of NAD^+^while varying prephenate concentrations. Inhibition kinetics were done by varying tyrosine concentrations 0, 8, 16, 24, 50, 100, 200 μM. Data analysis was conducted with Graphpad Prism 7® (GraphPad Software, San Diego, CA, USA) and Saturation and inhibition curves were fitted to the non-linear Michaelis-Menten equation.

### Phylogenetics

NCBI BLAST search engine was used to obtain bacterial prephenate dehydrogenase sequences (Altschul, Gish et al. 1990). *Aquifex aeolius* PDH (without ACT domain) (Sun, Shahinas et al. 2009) and *Bacillus anthracis* PDH (containing the ACT domain) (Shabalin, Gritsunov et al. 2020) served as the query sequences. The retrieved sequences were aligned by using ClustalW algorithm in MEGA X software with default parameters (Kumar, Stecher et al. 2018). A phylogenetic tree was constructed using the neighbour-joining method with 500 bootstrap replicates, the poisson substitution model, uniform rates, and a 95% cutoff for partial deletions. The resulting tree was visualized using the Interactive Tree of Live (iTOL) tool (Letunic and Bork 2016).

### Crystallization

ΔACT-BaPDH was crystallized using sitting-drop vapor-diffusion method. ΔACT-BaPDH (6 mg·mL^−1^) was supplemented with 5 mm NAD^+^ as needed. Aliquots of 0.2 µL of this solution were mixed with 0.2 µL of crystallization screening cocktail on 96-well, three-drop plates (Swissci, Neuheim, Switzerland) using a Mosquito crystallization robot (TTP Labtech, Hertfordshire, UK) and equilibrated against 1.5 m NaCl at 16 °C. Crystals were obtained in 20% w/v PEG3350, 0.2M Li citrate, and diffracted to ∼ 2.4 Å, speace group P1211. .

### Data collection, processing and structure determination

Crystals were harvested and vitrified in liquid nitrogen with cryoprotection by Paratone-N (Hampton Research, Aliso Viejo, CA, USA) or by slow dehydration (keeping the crystal over 1 m sodium chloride solution for 10 min). Diffraction data were collected at 100 K at the 19-BM beamline (19-BM, Lemont, IL, USA) at the Advanced Photon Source (UChicago Argonne LLC, Lemont, IL, USA).

Data reduction and scaling for all datasets were performed with HKL-3000 (Otwinowski and Minor 1997, Kutner, Shabalin et al. 2018). The structure of ΔACT-BaPDH in complex with NAD^+^ was solved by molecular replacement by HKL-3000 with the use of molrep (Minor, Cymborowski et al. 2006) and other programs from the ccp4package (Vagin and Teplyakov 2010). The structural model of full-length BaPDH (PDB ID:5UYY) was used as the template (Shabalin, Gritsunov et al. 2020). Automated model building was performed with Buccaneer (Winn, Ballard et al. 2011), followed by optimization of side-chain conformations with Fitmunk (Cowtan 2006), both integrated with HKL-3000. The structures were refined with HKL-3000 using state-of-the-art guidelines outlined elsewhere(Porebski, Cymborowski et al. 2016). refmac (Shabalin, Porebski et al. 2018) in restrained mode with automatic local NCS and hydrogen atoms in riding positions was used for the reciprocal-space refinement. Coot (Murshudov, Skubak et al. 2011) was used for the visualization of electron density maps and manual inspection and correction of the atomic models. TLS groups were introduced in the latest stages of refinement with the TLS Motion Determination Server (Emsley, Lohkamp et al. 2010). The TLS parameters were kept if confirmed by a significantly improved Rfree and the Hamilton R-factor ratio test (Painter and Merritt 2006) as implemented in HKL-3000. Structures were validated with a stand-alone version of Molprobity (Merritt 2012) and wwPDB validation tools (Chen, Arendall et al. 2010). The atomic coordinates and structure factors have been deposited in the PDB (PDB ID:7MQV). Structural figures (Figs 2-4) were prepared with the use of pymol (Schrodinger LLC, New York, NY, USA).

**Figure 1.**
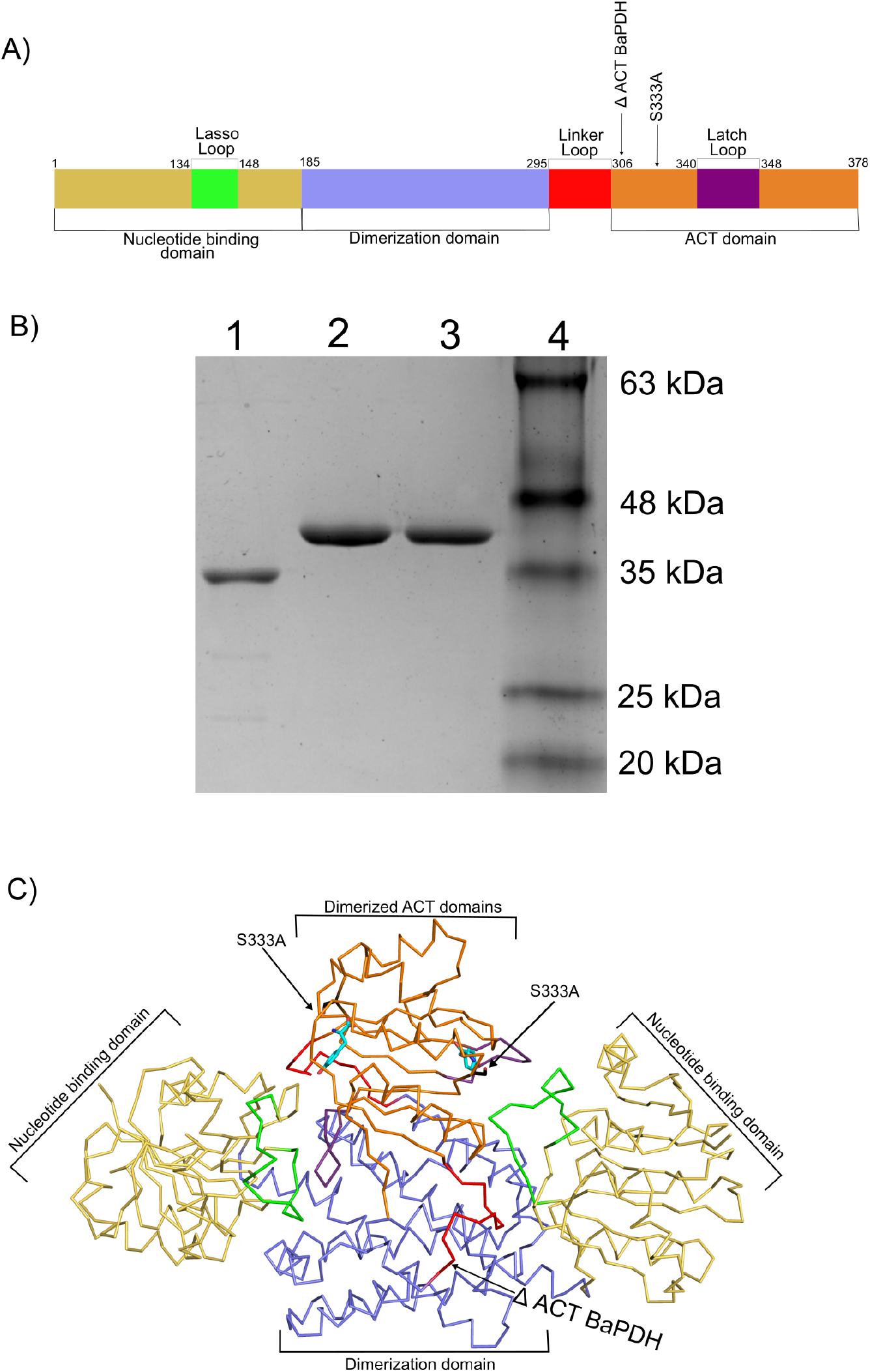
Components of full length and truncated BaPDH constructs. A) Diagram representation of domains and mutations on wild type and mutant BaPDH proteins. B) Gel of purified proteins for ΔACT-BaPDH(1), S333A(2),WT(3) and molecular weight ladder (4). C) Full length BaPDH structure (PDB id:6U60).

**Figure 2.**
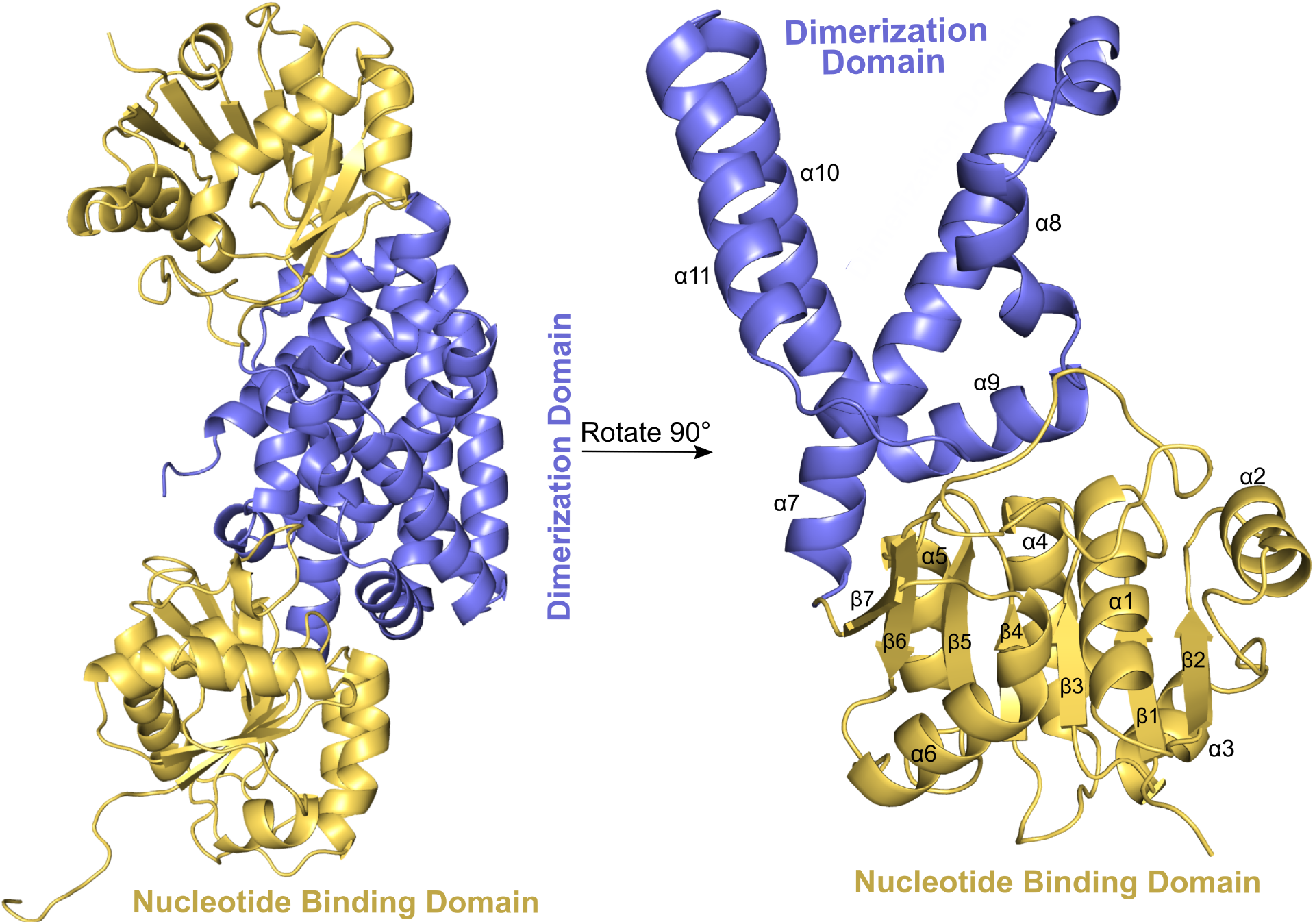
ΔACT-BaPDH (PDB:7MQV) dimer and a monomer. Dimerization domains are represented by blue color, whereas nucleotide binding domain golden. Secondary structure motifs were labeled on the monomer structure.

## Results

### Structural analysis of truncated and full length BaPDH constructs

PDH plays a central role in tyrosine biosynthesis by ctalyzing the conversion of prephenate to 4-hydroxyphenylpyruvate (Sun, Shahinas et al. 2009). A subset of PDHs contains an ACT domain, which allows these enzymes to be allosterically inhibited by tyrosine (Shabalin, Gritsunov et al. 2020). We therefore investigated the mechanistic and structural roles of the ACT domain, as well as the evolutionary relationships of ACT containing PDHs with other PDH variants.

To explore these aspects, we purified a truncated BaPDH construct (BaPDH 1-306) lacking the C-terminal ACT domain (ΔACT, residues 307-378), and crystallized it for structural studies (Figure 1 and 2, Crystal diffraction and refinement parameters are listed in Table 2.).The resulting structure revealed that the truncated protein forms a dimer with strong inter-helical interactions at the dimer interface. This observation of the dimeric organization mirrors that of the full-length ACT-containing BaPDH, which also exists as a dimer in solution, as confirmed by gel-filtration chromatography (Shabalin, Gritsunov et al. 2020). The overall structural topology of the dimeric ΔACT-BaPDH resembles a dumbbell, with the dimerization domain flanked by two outer catalytic domains, which is analogous to the weights on a dumbbell (Figure 2). The secondary structure of this truncated protein closely resembles both the full-length BaPDH and the *Aquifeux aeolicus* PDH (Aa-PDH) structure described elsewhere (Sun, Shahinas et al. 2009). Specifically, the ΔACT-BaPDH dimerization domain comprises a six-helical bundle, with each monomer contributing three helices: α7 (residues 186–219), α10 (252–275), and α11 (278–294). The N-terminal region forms a dinucleotide-binding domain reminiscent of the Rossmann fold, which constitutes the outer segments of the dumbbell topology. (Figure 2). The N-terminal region forms a dinucleotide-binding domain reminiscent of the Rossmann fold, which constitutes the outer segments of the dumbbell topology. This domain, spanning residues 1–185, primarily facilitates NAD^+^ binding and, to a lesser extent, prephenate binding. A detailed structural description of this region is available in previous studies (Sun, Shahinas et al. 2009, Shabalin, Gritsunov et al. 2020). A comparison of conformational features between the ACT containing and the truncated structures will be the focus of our analysis. To this end, we performed structural superimposition of monomers from the full-length BaPDH (Figure 3A) and the ΔACT-BaPDH (Figure 3B).

**Figure 3.**
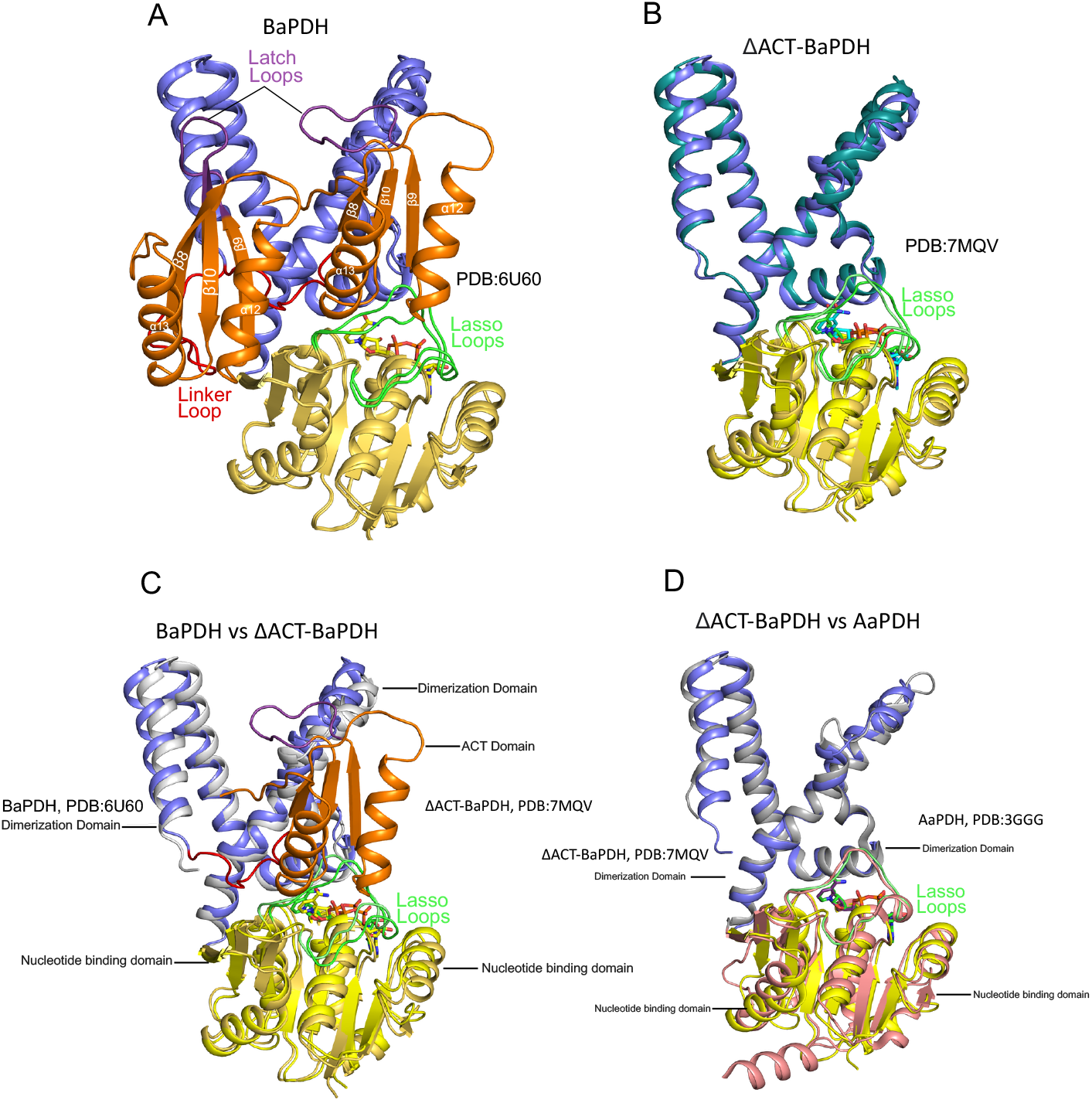
Superimposition of various PDH monomers from *A. aeolicus, B. anthracis*. A) BaPDH monomers. B) ΔACT-BaDPH monomers C) BaPDH vs ΔACT BaPDH D) *Aquifex* and ΔACT-BaPDH

The two monomers within the full-length BaPDH dimer exhibit distinct conformational differences, particularly at the active site and the ACT–prephenate binding interface. Notably, β-strands 8, 9, and 10 are shorter in one monomer, contributing to the formation of a larger loop region known as the Latch Loop(Shabalin, Gritsunov et al. 2020). This loop extends into the nucleotide-binding pocket of the active site, which notably lacks a bound NAD^+^ molecule. In contrast, the second monomer shows a more extended β-strands 8′, 9′, and 10′, resulting in a shorter Latch Loop region that adopts an alternate conformation which is oriented away from the NAD^+^ binding site. Interestingly, this site is occupied by a NAD^+^ molecule in the full-length protein. It is also important to note that full-length BaPDH crystalized without NAD^+^ cofactor shows a similar open confirmation of the lasso loop, even in the absence of cofactor. Additionally, the loop connecting the ACT domain to the protein is positioned within the active site, thus obstructing substrate access to the binding site. This structural insight guided the design of our ACT-truncated constructs. Two variants were successfully expressed and purifed and and thus were used in subsequent analysis: in one variant we retained the connecting loop, while it was excluded in the other variant. Although both constructs were expressed and purified, only the ΔACT-BaPDH variant containing the connecting loop was shown to be enzymatically active and therefore selected for further kinetic and structural studies.

Analysis of the BaPDH three dimensional protein structures was done to understand if the conformational change observed amongst the two molecules of the full-length ACT containing BaPDH is retained in this ΔACT-BaPDH construct. We first compared the two truncated ΔACT-BaPDH molecules in the dimer to identify conformational differences. They superposed very well, with an RMSD of 1.2Å (Figure 3 B) indicating no observable conformational difference between them. Further, the lasso loop is positioned within the active site to allow for the efficient binding of the substrate and cofactor NAD^+^ (Fig 4.3 B). In support of this we observed one molecule of NAD^+^ occupying each cofactor binding site. Since the full-length protein binds a single NAD^+^ molecule in the dimeric state, we compared the structure of individual molecules of the dimer to a ΔACT-BaPDH monomer to identigy conformational changes associated with NAD^+^ binding. The NAD^+^ bound monomer of the full-length protein superposed well (Figure 3 C), 1.6 Å RMSD, with the ΔACT-BaPDH molecule. In contrast, the lasso loop region, residues 134-148, of the NAD^+^ free molecule adopts an alternate conformation, that occluded the NAD^+^ binding site. This conformation of the lasso loop is stabilized by a set of H-bonding interactions with the latch loops. Residues of the lasso loop (chain A) M134, G136, L149, E151, form hydrogen bond interactions with G345, R343 of the latch loop (chain B). Since the truncated protein, ΔACT-BaPDH, does not have an ACT domain these interactions were not observed. This frees up both nucleotide binding sites of the truncated construct for NAD^+^ binding. These observations are indicative that conformational change with the lasso and latch loops governs NAD^+^ accessibility to the enzyme active site.

The effect of the interaction between the ACT domain, latch loop, and lasso loop extends into the nucleotide-binding domain in a domino-like fashion, resulting in an overall elongation of the BaPDH dimer by approximately 7 Å (Figure 4). In the full-length structure, residues H148, E151, and N152 of the nucleotide-binding domain (chain A) appear to form hydrogen bonds with residues D312, V313, and D315 of the ACT domain (chain B). The ACT domain interfaced tightly onto the nucleotide-binding domain via the lasso loop, which connects β-sheets 5 and 6. As a result, the lasso loop residues E151, A147, and K146 are displaced by 2–4 Å compared to the truncated structure. This lasso loop spans 10 amino acids and interacts with multiple secondary structural elements within the nucleotide-binding domain. These interactions contribute to an additional shift of 2–4 Å relative to the truncated construct. Overall, the ACT domain (chain B), positioned atop the BaPDH dimer, appears to induce significant conformational changes not only in the lasso loop but also across other regions of the nucleotide-binding domain (chain A).

**Figure 4.**
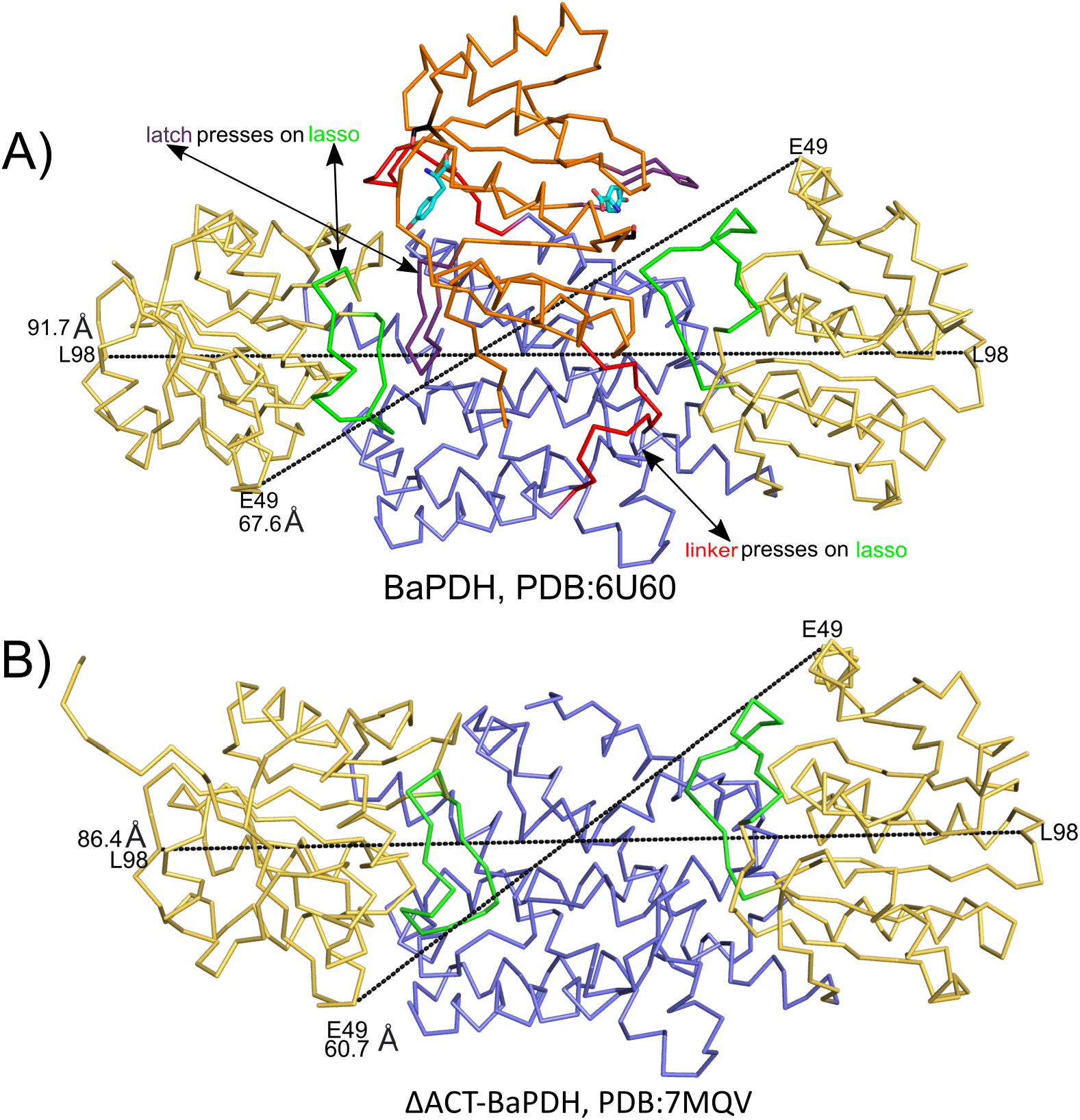
ACT domain on BaPDH causes the catalytic domain to extend outwards by 7 angstroms if compared to the truncated form. A) Dimer of full length PDH and measurements from end to end of the protein (in angstroms). B) Dimer of truncated BaPDH with measurements from end to end (in angstroms). Full-length BaPDH dimer appears to be longer by 5-7 angstroms compared to the truncated dimer. ACT domain that sits on top of the BaPDH dimer, presses onto the lasso and NTD domains, resulting in an extended dimer conformation.

Structural analysis revealed that the full-length BaPDH dimer is approximately 7 Å longer than the truncated construct (Figure 4A, B). This elongation is attributed to conformational shifts in the nucleotide-binding domains, driven by the presence of the ACT domain. The ACT domain sits atop the dimerization interface and interacts asymmetrically with only one monomer, resulting in the shortening of β-strands 8, 9, and 10 in that monomer. In contrast, the second monomer does not exhibit this structural distortion. These non-symmetrical interactions likely contribute to the allosteric regulatory mechanism affecting substrate binding. Specifically, the β8 strand of the ACT domain forms a stabilizing interaction with residue R235, which is involved in binding the carboxyl group of prephenate. Additionally, the linker loop connecting β8 positions residue P306 directly within the active site, obstructing prephenate binding. In the truncated construct, however, the linker loop is oriented away from the active site, allowing unobstructed access for substrate binding.

### Site-directed mutagenesis of S333A in the ACT domain

In order to investigate the structural and mechanistic basis of allosteric inhibition of BaPDH by tyrosine, we generated site-directed mutations in the tyrosine-binding site of the ACT domain. Residue S333 was specifically targeted for mutagenesis, as it resides within the ACT domain and hydrogen bond with bound tyrosine. We attempted to crystallize the S333A variant under tyrosine-supplemented conditions to elucidate conformational changes associated with tyrosine-mediated allosteric inhibition. Although crystals of the S333A variant were successfully obtained, they did not yield diffraction data of sufficient quality for structural determination. Nevertheless, we conducted detailed enzymatic assays to evaluate the functional role of S333 in tyrosine binding, as described in subsequent sections.

### Enzyme kinetics and analysis of tyrosine as an inhibitor of BaPDH

Although the ACT domain of PDH has been implicated in allosteric regulation of enzymatic activity, its precise role remains to be fully established. Based on the three-dimensional structure of the enzyme and the positioning of active site residues, we generated an ACT-truncated construct and site-specific variants targeting residue S333 (Figure 5). Since S333 is located within the ACT domain and forms a hydrogen bond with tyrosine, we hypothesized that mutating this residue would alleviate tyrosine-mediated inhibition. Similarly, we proposed that removing the ACT domain would eliminate allosteric regulation by tyrosine.

**Figure 3.**
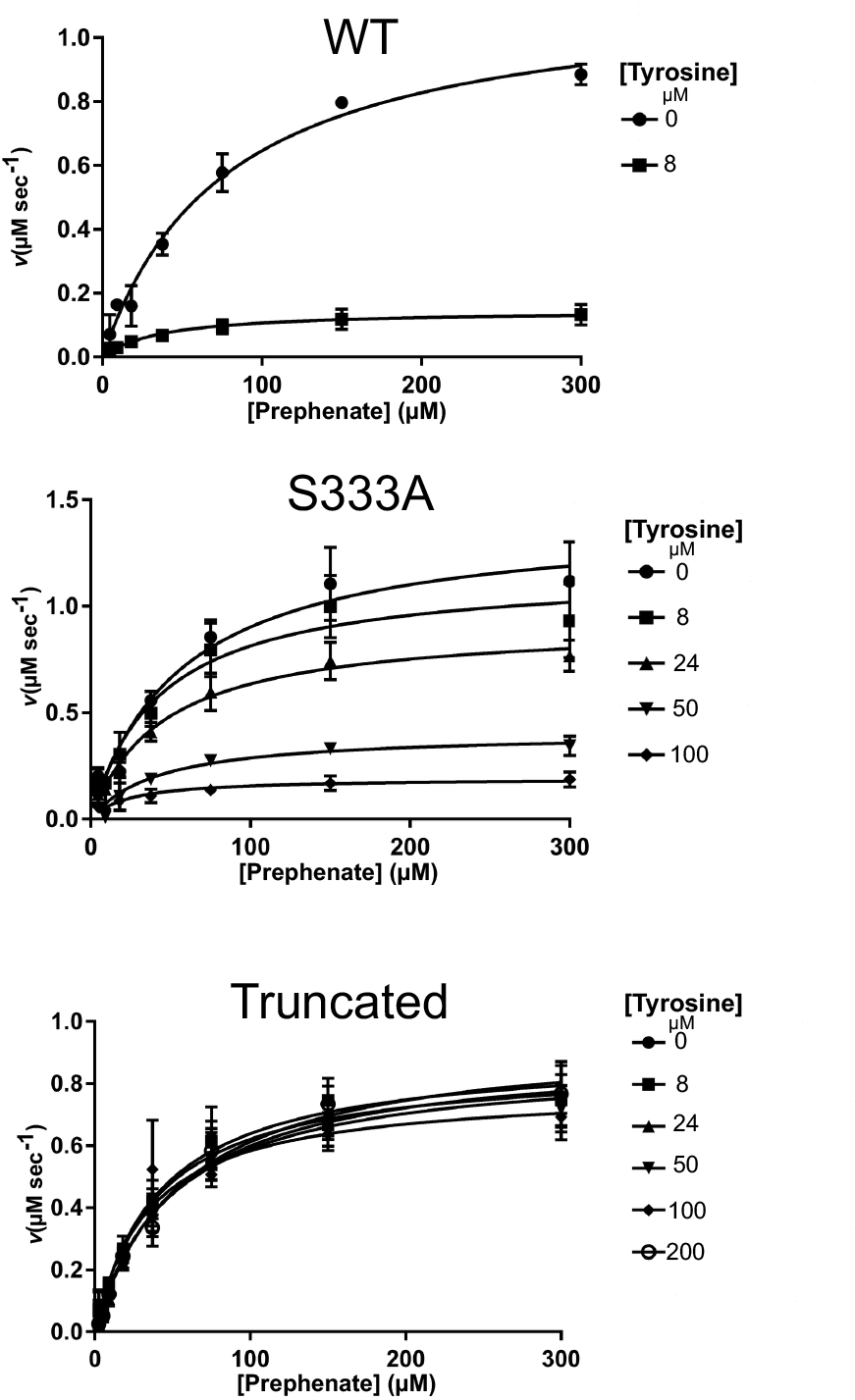
Enzyme kinetics analysis for WT, SS333A and the ΔACT-BaPDH construct. Tyrosine inhibition analysis was conducted with WT (top), S333A mutant (middle) and the truncated construct (lower) saturation plots.

**Table 1:**
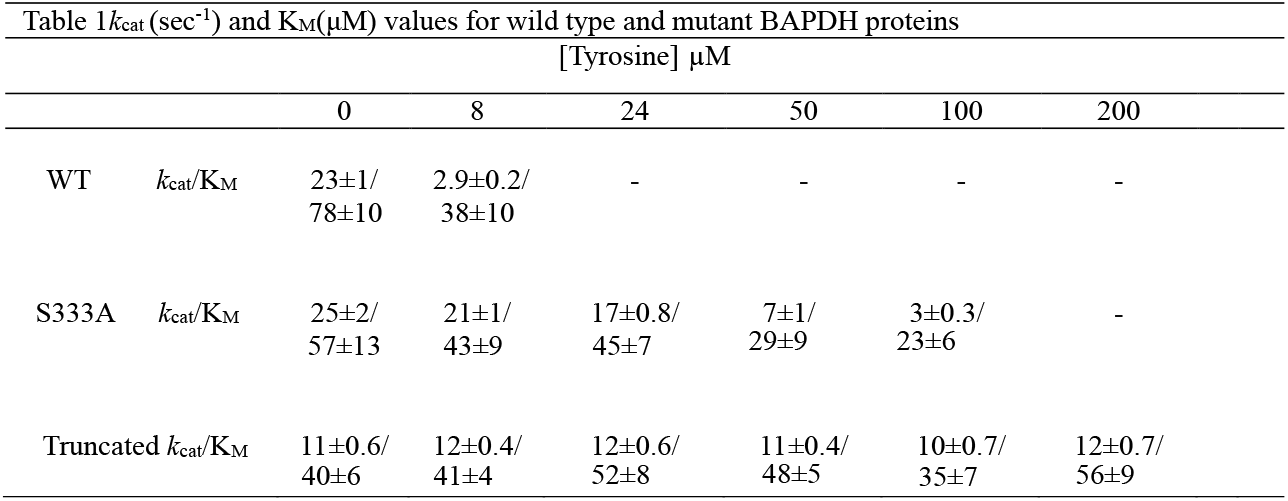
Kinetic parameters for the WT and PDH variants with varing concentrations of tyrosine.

To test these hypotheses, saturation plots were generated for both the truncated variant and the S333A mutant by varying prephenate concentrations below and above the established KM, while maintaining a fixed concentration of tyrosine. Enzymatic assays were repeated across a range of tyrosine concentrations, with the highest concentration being 25 times greater than that required for complete inhibition of the full-length enzyme. Saturation kinetics were used to determine the Michaelis constants and turnover rates for ΔACT-BaPDH, S333A, and full-length wild-type BaPDH (WT-BaPDH), as summarized in Figure 5.

WT-BaPDH activity was fully inhibited at 8 μM tyrosine, with an inhibition constant (K_I_) of 1.5 ± 0.2 µM. In contrast, the S333A variant required 100 μM tyrosine for inhibition, with a K_I_ of 32 ± 2 µM. These values are significantly higher than that for the wild-type enzyme. Much more importantly, the truncated ΔACT-BaPDH construct showed no sensitivity to tyrosine, with no inhibition observed even at concentrations up to 200 μM.

### Phylogenetic analysis of PDH-ACT proteins distribution in bacteria

It is expected that the subset of PDHs that contain the ACT domain are subjected to allosteric regulation. Therefore, we investigated the evolutionary trajectories underlying ACT domain acquisition in PDH proteins by examining the distribution of ACT containing PDHS amongst different organisms and the environment in which these organisms proliferate. Sequences of ACT-containing PDHs were retrieved using the NCBI BLAST search engine (Altschul, Gish et al. 1990), with the BaPDH amino acid sequence (Mmdb_id: 147981) as the query and a stringent E-value cutoff of < 1e–50.

To ensure this is a representative PDH dataset, we also included non-ACT-containing PDHs, such as the AaPDH sequence. Each retrieved protein was considered a functional PDH based on the presence of conserved catalytic and substrate-binding residues, as illustrated in the WebLogo representation (Figure 6). Key active site residues identified include S109, H132, S198, S199, S202, and R235, which are known to be essential for substrate binding (Sun, Shahinas et al. 2009). Additionally, residue W244 was found to confer substrate specificity for prephenate over arogenate (Legrand, Dumas et al. 2006).

**Figure 4.**
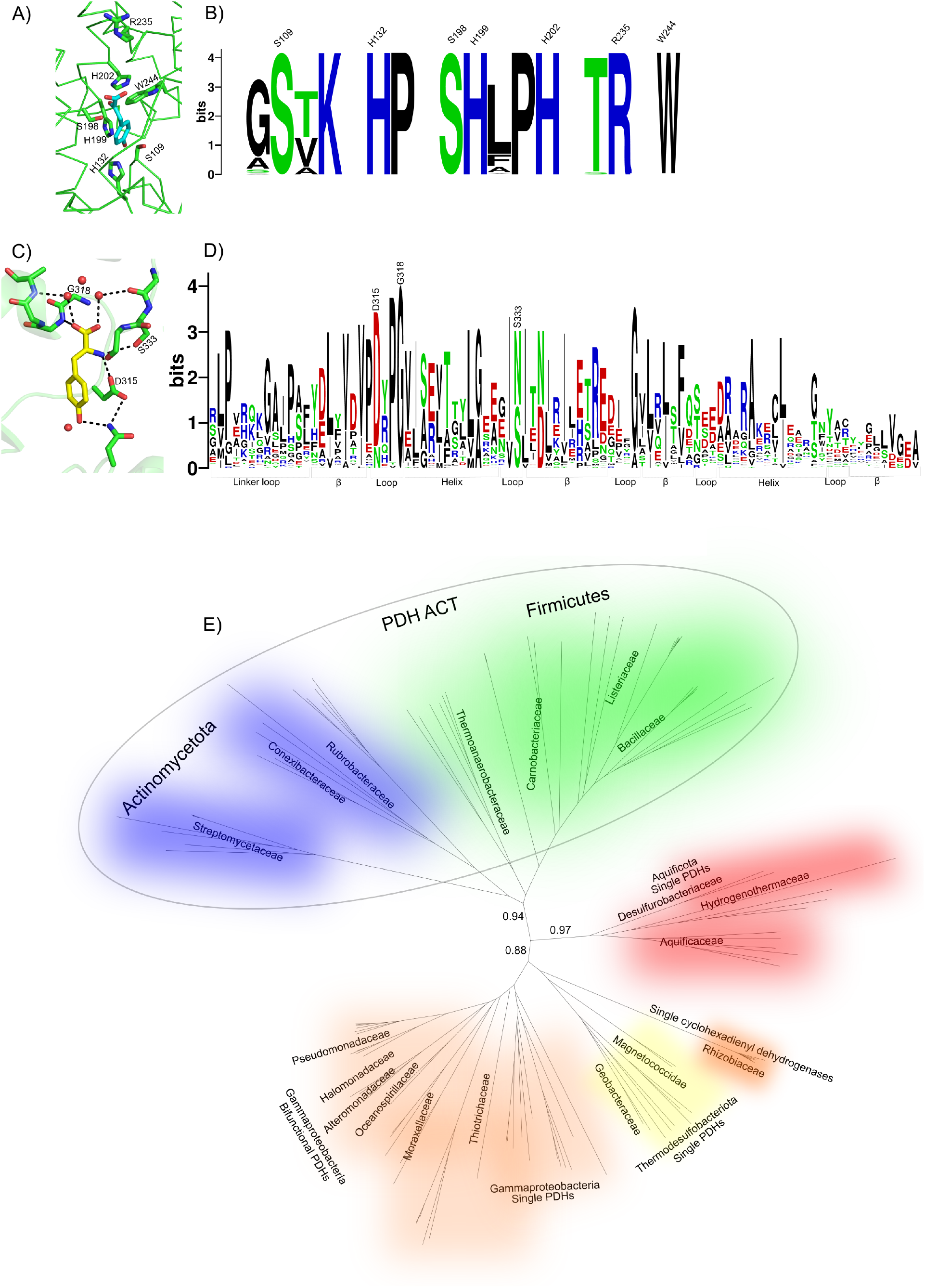
Phylogenetic relationships of PDHs between different groups of bacteria. A) and B) Active site residue conservation among PDHs subjected to the phylogenetic analysis. C) and D) Several ACT domain tyrosine binding residues such as G318, D315, S333 are conserved or replaced by similar residues. E) Phylogenetic analysis of PDH sequences from bacteria reveals *Actinomycetota* and *firmicute* species to possess PDH-ACT domain fusion. The other PDH group from *aquificota* have only PDHs with no-ACT fusion. Several *Alphaproteobacteria* groups such as rhizobiaceae, magnetococcidae also have PDHs without the ACT-fusion. Similarly, single gammaproteobacteria PDHs were identified in Thiotrichaceae. Other type of PDHs exist on a bifunctional peptide fused to chorismate mutase. These proteins are found in gammaproteobacteria groups.

The presence of the ACT domain was confirmed through secondary structure prediction and modeling. Analysis using established methods (Montgomerie, Sundararaj et al. 2006) revealed that putative ACT domains exhibit the characteristic βαββαβ fold. Within these domains, tyrosine-binding residues specific to the ACT domain were identified. Notably, residues D315 and S333, which interact with tyrosine via their side chains, are replaced by asparagine (N) residues in Actinomycetota species, suggesting a conserved functional adaptation across this group.

Phylogenetic reconstruction of the retrieved PDH sequences revealed their distribution into two distinct clusters. ACT-containing PDHs formed a separate clade from non-ACT PDHs, which are known to be competitively regulated. Notably, ACT-containing PDHs were predominantly found in species belonging to the Firmicutes and Actinomycetota phyla. Within Firmicutes, these PDHs were distributed across families such as *Thermoanaerobacteraceae, Carnobacteriaceae, Listeriaceae*, and *Bacillaceae*. Similarly, Actinomycetota PDHs were found among *Streptomycetaceae, Conexibacteraceae*, and *Rubrobacteraceae*. This distinct phylogenetic clustering suggests that the acquisition of the ACT domain likely occurred in a common ancestor of Firmicutes and Actinomycetota within the Terrabacteria group, representing a significant evolutionary event in the regulation of PDH enzymes.

## Discussion

PDH plays an important role in the highly regulated biosynthesis of tyrosine and downstream aromatic compounds. PDHs have been shown to be competitively regulated by tyrosine in a large subset of microorganisms (Sun, Shahinas et al. 2009). However, some PDHs contain an ACT domain and the regulation of these proteins has only recently became apparent (Shabalin, Gritsunov et al. 2020). In this study we undertake a structural and biochemical approaches to investigate the mechanism of regulation of the ACT containing PDHs and also the phylogenetic distribution to understand the biological importance for the evolution of the ACT mediated regulation of PDHs.

The three dimensional crystal structure of the ACT truncated PDH from *Bacillus anthracis* was determined and compared to the full-length ACT containing protein as well as to the competitively regulated *Aquifex aeolicus* PDH. The full-length protein is a dimer with each monomer adopting distinct conformations especially in regions that are contributing to the enzyme active site. The most dramatic conformational differences are observed in the Lasso, Latch and Linker loops which are thought to be the regions that regulate substrate binding. In order to understand the conformational and mechanistic roles of these regions in modulating substrate binding and thus enzyme function we prepared an ACT trunctated construct of BaPDH and conducted somparative structural and biochemical analyses. In contrast to the full-length protein, both monomers of the truncated protein adopt similar conformations including their Lasso and Latch loop regions. This is in contrast, the ligand free full-length BaPDH monomer which adopts distinct conformations from that of the truncated protein. We postulated that these conformational differences are directly attributed to the allosteric mechanism of regulation of BAPDH by the ACT domain. This is further supported by the fact that tyrosine binding to the ACT domain results in conformational changes in the three loop regions mentioned above which directly perturbs the binding of both the NAD^+^ cofactor and prephenate. Particularly it is quite evident that with tyrosine binding, the conformation of the Lasso loop becomes positioned directly in the NAD^+^ cofactor binding site thus occluded its binding. Similarly, the linker loop moves into the substrate binding site upon tyrosine binding to the ACT domain. The most distinct conformational change occurs at the substrate binding pocket in which Pro306 becomes directly positioned in the prephenate binding site. We reasoned that this multiple conformational changes, blocking both the cofactor and the substrate binding sites ensure strict control on the regulation of PDH activity. This level of allosteric control on PDH activity is much more powerful than competitive inhibition, whch can be allivated with increasing concentration of prephenate.

Enzymatic assay of the ACT truncated protein shows that it retains full PDH activity and has similar kinetic properties to that of the wild-type protein thus indicating that the removal of the ACT did not impart any undesirable structural/conformational changes to the protein. Consistent with the expected regulatory role of ACT domain, the truncated variant is insensitivie to tyrosine. These findings indicate that the ACT domain is modular and separable.

Intruigingly, a comparison of the substrate binding site between BaPDH and AaPDH indicated that all the functionally important amino acids are conserved. Since AaPDH is competitively inhibited by tyrosine, we investigated if higher concentration of tyrosine can also affect truncated BaPDH activity. However, high concentration of tyrosine did not impart any inhibitor effect on the protein. This was somewhat surprising to us since the active site architecture and amino acid composition are very similar in both truncated BaPDH and AaPDH. We expected that the open active site of the truncated BaPDH will allow for free access to tyrosine to compete with prephenate binding. We reasoned that the inability to competitively inhibit the truncated BaPDH is attributed to main-chain conformations within the substrate binding site. A detailed analysis of the main-chain conformation and sequence variation between AaPDH and truncated BaPDH revealed a region in the prephenate binding site that is known to confer tyrosine specificity to PDH. Mutational analysis of these main chain residues in the truncated protein, expecially a AGG motif, did not enable competitive tyrosine inhibition. We therefore concluded that additional structural and conformational elements away from the binding site of BaPDH are required to confer tyrosine binding as such will be the focus of future studies.

The structural and biochemical analyses allow us to put forward a model in which high tyrosine concentration induces allosteric inhibition of the ACT-contining BaPDH. That is, tyrosine binding to the ACT domain triggers a conformational shift in the lasso loop region. This movement repositions the loop into the substrate and cofactor binding site, thereby preventing either ligand from docking onto the enzyme. Under conditions of low intracellular tyrosine concentration the ACT domain is unbound, leaving the substrate and cofactor binding site fully accessible. Therefore, PDH activity will proceeds because the enzyme remains in its active state.

Based on our phylogenetic analysis, we observed that many organisms containing the ACT-PDH colonize environment with limited resources and some are in extreme conditions. One can speculate then that the acquisition of the ACT domain in these PDH allows the enzyme to be efficient in the utilization/partitioning of chorismate for aromatic compound biosynthesis. This is supported by the colonization of a large reporitoire of different strains of Firmicutes organisms in adverse and extreme environments including hotsprings, salt lakes and salterns where carbon nutrient is limited. Under these growth conditions, organisms need to conserv resources and have strict control over how they are used for the biosynthesis of essential metabolites. In addition to elucidating the evolutionary advantage conferred by ACT domain acquisition, our mechanistic insights into ACT-mediated regulation of prephenate dehydrogenases (PDHs) provides promising opportunities for therapeutic intervention. Specifically, inhibition of *Bacillus anthracis* PDH (BaPDH) may offer a novel strategy to suppress the growth of this and other ACT-containing pathogenic bacteria. Previous studies have shown that tyrosine limitation, either through genetic disruption of PDH or nutrient deprivation, impairs the ability of *Bacillus cereus* to sporulate and form biofilms (Champney and Jensen 1970, Kiley and Stanley-Wall 2010, Huijboom, Tempelaars et al. 2023). Furthermore, tyrosine serves as a critical precursor in the biosynthesis of numerous pharmacologically relevant compounds, including neurotransmitters and hormones (Hagel and Facchini 2013). Thus, engineering a constitutively active PDH variant lacking the ACT domain represents a promising approach for enhancing tyrosine production in industrial biotechnology applications.

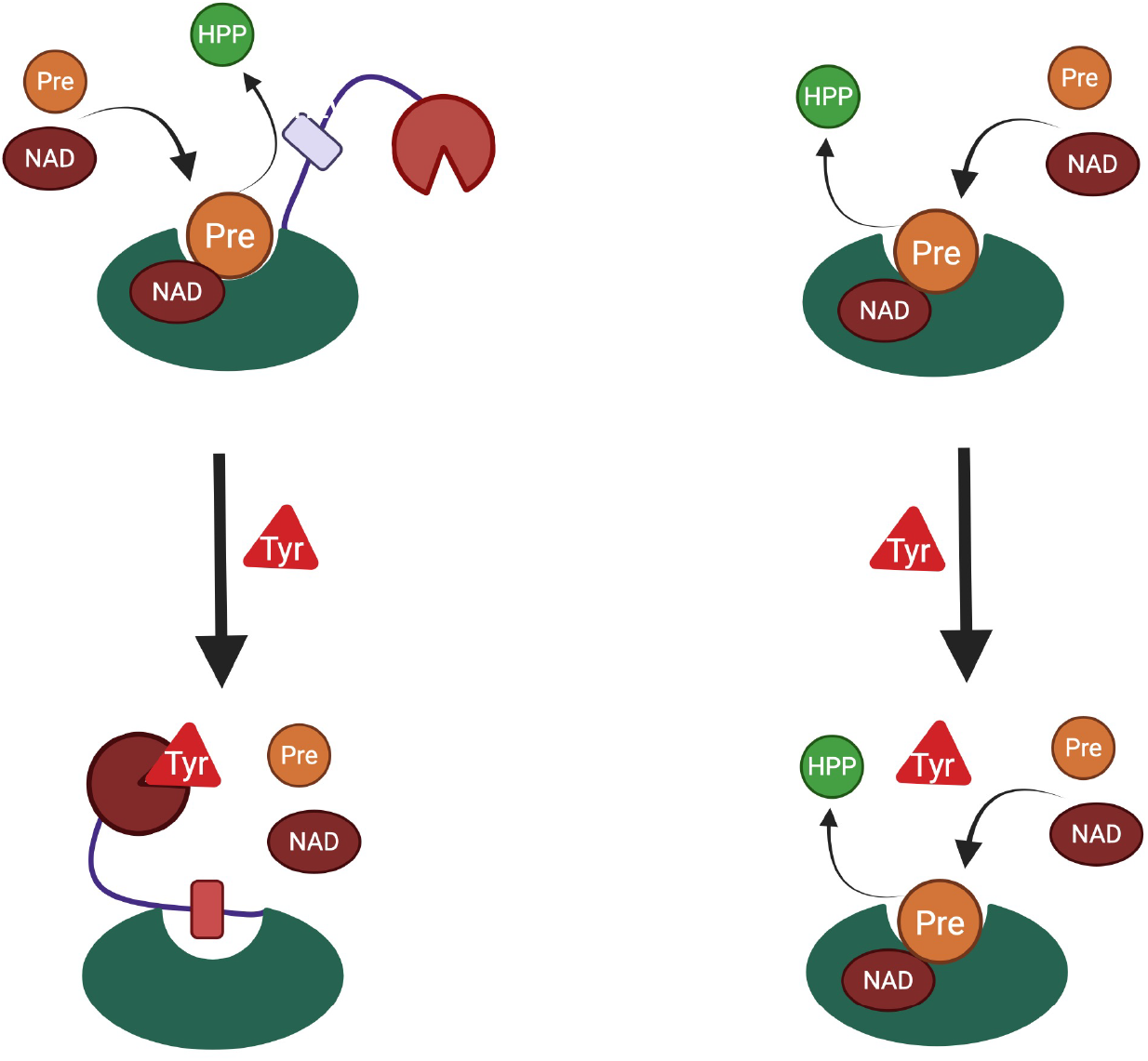
Model for allosteric inhibition of BaPDH by tyrosine. **Left panel:** ACT-BaPDH, elevated tyrosine level leads to tyrosine binding to the ACT domain and reposition the lasso loop into the substrate binding site. This conformation prevents the binding of NAD+ and prephenate to the enzyme and thus keeps it in an inactive state. **Right panel:**Truncation of the ACT domain from BaPDH results in a tyrosine insensitive enzyme even at artificially high concentration.

**Table 2:** Data collection and structure refinement statistics.

| **Data collection** |  |
| --- | --- |
| Resolution (Å) | 50.00 - 2.40 (2.44 - 2.40) |
| Wavelength (Å) | 0.97929 |
| Space group | P21 |
| a, b, c (Å) | 73.77, 118.88, 75.56 |
| $\alpha$, $\beta$, $\gamma$ (°) | 90, 92.39, 90 |
| Completeness (%) | 99.7 (98.2) |
| Reflections used | 158602 |
| $\langle I \rangle / \langle \text{Sigma } I \rangle$ | 9.4 (1.1) |
| Redundancy | 3.1 (2.9) |
| Rmerge | 0.120 (0.868) |
| Rpim | 0.080 (0.592) |
| CC1/2 last shell |  |
| Wilson B factor (Å<sup>2</sup>) | 41.2 |
| **Refinement** |  |
| Rwork / Rfree | 0.168 / 0.225 |
| Resolution (Å) | 49.12 - 2.40 |
| Reflections all | 44898 |
| Reflections for R<sub>free</sub> | 2131, 4.7% |
| Bond lengths rmsd (Å) | 0.007 |
| Bond angles rmsd (°) | 1.45 |
| Mean B value (Å<sup>2</sup>) | 47 |
| Number of protein atoms | 9095 |
| Mean B value for protein atoms (Å<sup>2</sup>) | 47 |
| Number of water atoms (expected) | 279 (595) |
| Mean B value for water atoms (Å<sup>2</sup>) | 39 |
| Number of ligands/ions atoms | 177 |
| Mean B value for ligands/ions atoms (Å<sup>2</sup>) | 37 |
| Clashscore | 3.37 |
| Clashscore percentile (100) | 88.8 |
| Rotamer outliers (<1%) | 2.54 |
| Ramachandran outliers (<0.2%) | 0.00 |
| Ramachandran favored (>98%) | 96.70 |
| Residues with bad bonds (<0%) | 0.43 |
| Residues with bad angles (<0.1%) | 1.55 |
| MolProbity score | 1.64 |

## Acknowledgements

The presented work was supported by federal funds from the National Institute of Allergy and Infectious Diseases, National Institutes of Health, Department of Health and Human Services under contracts HHSN272201200026C and HHSN272201700060C; National Institute of General Medical Sciences under grant numbers GM118619 and GM117325; and by the Natural Sciences and Engineering Research Council of Canada Discovery Grant (NSERC-DG RGPIN-2020-06052). Results shown in this report are derived from work performed at Argonne National Laboratory, operated by UChicago Argonne, LLC, for the U.S. Department of Energy, Office of Biological and Environmental Research under contract DE-AC02-06CH11357. Use of the LS-CAT Sector 21 was supported by the Michigan Economic Development Corporation and the Michigan Technology Tri-Corridor (Grant 085P1000817).

